# Potentiation of Long-Acting β_2_-Agonist and Glucocorticoid Responses in Human Airway Epithelial Cells by Modulation of Intracellular cAMP

**DOI:** 10.1101/2020.11.09.375089

**Authors:** Yechan Kim, Vincent Hou, Ryan D. Huff, Jennifer A. Aguiar, Spencer Revill, Nicholas Tiessen, Quynh Cao, Matthew S. Miller, Mark D. Inman, Kjetil Ask, Andrew C. Doxey, Jeremy A. Hirota

## Abstract

**Introduction:** Over 300 million people in the world live with asthma, resulting in 500,000 annual global deaths with future increases expected. It is estimated that around 50-80% of asthma exacerbations are due to viral infections. Currently, a combination of long-acting beta agonists (LABA) for bronchodilation and glucocorticoids (GCS) to control lung inflammation represent the dominant strategy for the management of asthma, however it is still sub-optimal in 35-50% of moderate-severe asthmatics resulting in persistent lung inflammation, impairment of lung function, and risk of mortality. Mechanistically, LABA/GCS combination therapy results in synergistic efficacy mediated by intracellular cyclic adenosine monophosphate (cAMP).

**Hypothesis:** Increasing intracellular cAMP during LABA/GCS combination therapy via inhibiting phosphodiesterase 4 (PDE4) and/or blocking export of cAMP by ATP Binding Cassette Transporter C4 (ABCC4), will potentiate anti-inflammatory responses of mainstay LABA/GCS therapy.

**Methods:** Expression and localization experiments were performed using *in situ* hybridization and immunohistochemistry in human lung tissue from healthy subjects, while confirmatory transcript and protein expression analyses were performed in primary human airway epithelial cells and cell lines. Intervention experiments were performed on the human airway epithelial cell line, HBEC-6KT, by pre-treatment with combinations of LABA/GCS with PDE4 and/or ABCC4 inhibitors followed by Poly I:C or imiquimod challenge as a model for viral stimuli. Cytokine readouts for IL-6, IL-8, CXCL10/IP-10, and CCL5/RANTES were quantified by ELISA.

**Results:** Using archived human lung and human airway epithelial cells, ABCC4 gene and protein expression were confirmed *in vitro* and *in situ*. LABA/GCS attenuation of Poly I:C or imiquimod-induced IL-6 and IL-8 was potentiated with ABCC4 and PDE4 inhibition, which was greater when ABCC4 and PDE4 inhibition was combined. Modulation of cAMP levels had no impact on LABA/GCS modulation of Poly I:C-induced CXCL10/IP-10 or CCL5/RANTES.

**Conclusion:** Modulation of intracellular cAMP levels by PDE4 or ABCC4 inhibition is able to potentiate LABA/GCS efficacy in human airway epithelial cells challenged with viral stimuli. The data suggests further exploration of the value of adding cAMP modulators to mainstay LABA/GCS therapy in asthma for potentiated anti-inflammatory efficacy.

## INTRODUCTION

It is estimated that there are over 300 million people in the world with asthma, resulting in 500,000 annual global deaths (1). The Global Initiative for Asthma (GINA) defines asthma as a chronic inflammatory disease defined by history of respiratory symptoms such as wheeze, shortness of breath, chest tightness, and variable cough and expiratory airflow limitation (2). Key features of asthma include airway hyperresponsiveness, reversible bronchoconstriction, airway wall thickening, increased mucus production, and airway inflammation. Despite medications available to control symptoms, up to 35-50% of moderate-severe asthmatics experience sub-optimal symptom management, resulting in persistent lung inflammation, impairment of lung function, and increased risk of mortality (3–7). These individuals are more susceptible to asthma exacerbations that may be triggered by exercise, exposure to allergens or irritants, or respiratory infections (8–10). Approximately 80% of exacerbations are associated with respiratory tract viral infections including but not limited to: rhinovirus, influenza virus, adenovirus, and respiratory syncytial virus (11). Viral infections lead to inflammation of the lungs, via recruitment and productions of neutrophils, eosinophils, CD4^+^ cells, CD8^+^ cells, and mast cells through increased expression and secretion of IL-6, IL-8, CXCL10/IP-10, CCL5/RANTES, and other cytokines (12).

GINA guidelines outline evidence-based, stepwise symptom management strategies focused on control and risk reduction. Inhaled glucocorticoids (GCS) and long-acting β_2_ agonists (LABA) formulations form the basis for asthma control, decreasing lung inflammation and relieving bronchoconstriction (13). The mechanism of action for GCS is complex, involving both transrepression and transactivation processes that regulate gene expression networks important in immune responses (13,14). Inflammatory gene expression is induced by transcription factors such as Nuclear Factor κ-light-chain-enhancer of Activated B-Cells (NF-κB), and Activator Protein-1 (AP-1). GCS bind glucocorticoid receptors (GR) in the cytoplasm, resulting in dimerization, and translocation to the cell nucleus where they bind to glucocorticoid response elements (GREs), which in turn increase transcription of anti-inflammatory proteins or suppress transcription of pro-inflammatory genes (13,15).

LABAs elicit their effects by increasing intracellular cyclic adenosine monophosphate (cAMP) through the activation of adenylyl cyclase, which catalyzes the conversion of adenosine triphosphate (ATP) into cAMP. LABAs are key for mediating smooth muscle relaxation by elevating cAMP to enable protein kinase A (PKA) phosphorylation and regulation of proteins involved in muscle tone (16). Moreover, elevations of cAMP have been shown to inhibit calcium ion release from intracellular stores, reduction of membrane calcium ion entry, and sequestration of intracellular calcium ion, leading to relaxation of the airway smooth muscle (16). Importantly, elevations in cAMP in human airway epithelial cells may be related to induction of pro-inflammatory gene expression and cytokine production (17,18).

LABA/GCS have additive and synergistic effects on the regulation of anti-inflammatory and bronchoprotective genes. LABA/GCS *additively* increase the induction of the anti-inflammatory gene dual-specificity phosphatase (DUSP1), an inhibitor of MAPK(19). The MAPK pathway plays a key role in the transduction of extracellular signals to cellular response, including inflammation (20–23). *Synergistic* interactions of LABA/GCS has also been observed with regulator of G-protein signalling, RGS2, a bronchoprotective gene (24,25). RGS2 inhibits Gq protein, a G protein subunit that activates phospholipase C, which downstream, contributes to airway smooth muscle contraction. The interaction between LABA and GCS is hypothesized to involve PKA signalling interacting with promoter regions that are also influenced by GCS via GR binding to GREs (26). It follows that promoting PKA signalling through alternate mechanisms that elevate intracellular cAMP may render LABA/GCS more efficacious.

Intracellular cAMP levels are tightly regulated by enzymes and transporters (27). The PDE superfamily consists of enzymes that break phosphodiester bonds and are responsible for the hydrolysis of cAMP and cyclic guanosine monophosphate (cGMP) into their respective inactive form, AMP and GMP (28). PDE’s are sub-classified into 11 primary families with each one having differential preferences for cAMP or cGMP. PDE4, in particular, is highly relevant for chronic inflammatory airway diseases as it is the major regulator of cAMP levels in inflammatory cells (29,30). PDE4 has 4 different isoforms (A, B, C, D) with different localization in the cell. Working in partnership with PDE4 are transport mechanisms present in the cell membrane. The superfamily of ABC transporters uses ATP for the transport of substrates across membranes. For instance, ABCC4 can transport cAMP, uric acid, prostaglandins, and leukotrienes (31), and is expressed in the prostate, liver, testis, ovary, kidney, and the lungs (27). ABCC4 expression was first demonstrated in human airway epithelial cells for their ability to transport prostaglandins (32). Our group was able to demonstrate that ABCC4 can also transport cAMP and urate in human airway epithelial cells (33) and that inhibition of ABCC4 had functional consequences on cystic fibrosis transmembrane conductance regulator (CFTR) activity and LABA/GCS anti-inflammatory efficacy (34,35). Fine tuning of intracellular cAMP levels with interventions targeting enzymatic degradation and/or extracellular transport may have benefits for optimized management of asthma symptoms (36).

As current GINA guidelines recommend LABA/GCS as the primary method of asthma symptom management, add-on therapies that can optimize treatment efficacy in difficult to control patients may be of immediate relevance and value. The demonstrated interaction between elevated cAMP levels induced by LABA and anti-inflammatory gene expression by GCS, suggest that modulation of intracellular cAMP levels by additional mechanisms may provide benefit. In this study, we hypothesize that elevating intracellular cAMP levels via ABCC4 inhibition and/or PDE4 inhibition may increase LABA/GCS anti-inflammatory efficacy in human airway epithelial cells.

## MATERIALS AND METHODS

### Reagents

Formoterol and budesonide (Cayman Chemicals, Michigan, USA) were used as clinically approved LABA/GCS compounds. Ceefourin-1 [CF1] (Abcam, Massachusetts, USA) was used for ABCC4 inhibition (37). Roflumilast [RF], rolipram [RP] (Cayman Chemicals, Michigan, USA) and cilomilast [CM] (AdooQ Bioscience, California, USA) were used for PDE4 inhibition. Polyinosinic: polycytidylic acid [Poly I:C], a TLR3 agonist, and imiquimod, a TLR7 agonist were used as viral mimics (InvivoGen, California, USA).

### Human Airway Epithelial Cell Culture

*In vitro* experiments were performed using a human bronchial epithelial cell line (HBEC-6KT) derived from a healthy non-smoker by expression of human telomerase reverse transcriptase and cyclin-dependent kinase 4 grown under submerged monolayer conditions (38), which we have previously validated for innate immune responses (39–41). Cells were cultured in keratinocyte serum-free growth medium supplemented with 0.8ng/mL recombinant human epidermal growth factor (EGF), and 50 μg/mL Bovine Pituitary Extract and 1% Pen/Strep (ThermoFisher, Mississauga, Canada) at 37°C, 5% CO_2_, in a humidified incubator. Another human bronchial epithelial cell line (Calu-3) was obtained from ATCC (Manassas, Virginia, USA). Cells were grown in Minimum Essential Medium Eagle with 10mM HEPES (Sigma Aldrich, Oakville, Canada), 20% fetal bovine serum, and 1% Pen/Strep. Calu-3 cells were grown under submerged monolayer and fed in a two-day feeding cycle(42). Primary human airway epithelial cells were collected from clinical subjects by means of bronchial brushing. The brushings were washed with sterile PBS and the cells were collected and transferred to T-25 cm^2^ cell culture flasks. The cells were grown under submerged monolayer conditions as above in a two-day feeding cycle with PnemaCult™ Ex-Plus media supplemented according to manufacturer’s recommendations (Stemcell Technologies, Vancouver, Canada).

### Histology, digital slide scanning and microscopy

*In situ* hybridization and immunohistochemistry was performed using a Leica Bond Rx autostainer with instrument and application specific reagent kits (Richmond Hill, Ontario). Following selection of healthy primary lung tissues (n=10), four-micron thick sections were stained using RNAscope™ probes (*in situ* hybridization - RNAscope^®^ LS 2.5 Probe-Hs-ABCC4, California, USA) or antibody (immunohistochemistry - ABCC4 (M41-10 - Ab15602) primary antibody – Abcam, Toronto, Canada) following directions supplied with the Leica Bond reagent kits. For immunohistochemistry, heat-induced antigen retrieval in citrate buffer was performed at pH 6 with primary antibody diluted at 1:50. Slides underwent digital slide scanning using an Olympus VS120-L100 Virtual Slide System at 40X magnification with VS-ASW-L100 V2.9 software and a VC50 colour camera (Richmond Hill, Ontario). Image acquisition and formatting was performed using Halo Software (Indica Labs, New Mexico, USA).

### Gene Expression Dataset Curation, and Normalization

Public microarray experiments using Affymetrix chips (HG-U133 Plus 2, HuEx-1.0-st-v1 and HuGene-1.0-st-v1) on bronchial brushing samples of human airway epithelial cells from healthy individuals were selected from the NCBI Gene Expression Omnibus (GEO) database.

For all dataset samples, raw intensity values and annotation data were downloaded with the R statistical language (version 3.6.1; R Core Team, 2019) using the GEOquery R package (version 2.52.0) (43). Probe definition files were downloaded from the Brainarray database (version 24) (44). To obtain processed microarray gene expression values unaffected by probe CG compositional biases, the Single Channel Array Normalization (SCAN) method was used via the SCAN.UPC R package (version 2.26.0) (45) using annotation data from the Bioconductor project (version 3.9) (46). All log_2_-transformed gene expression data were unified into a single dataset, and only genes detected in all three platforms were kept for subsequent analyses. In order to reduce differential expression bias due to sex chromosome genes, only autosomal genes were included for subsequent analysis. Correction of experiment-specific batch effects was performed using the ComBat method (47) implemented in the sva R package (version 3.32.1) (48), with disease status and sex supplied as covariates. Following batch correction, all data underwent Z-Score transformation to set the mean of all samples to zero and replace expression values with a measure of variance from the mean (49). Datasets were filtered to include only healthy individuals, and these healthy samples were further filtered by removing former or current smokers. This resulted in a total of 616 samples from healthy subjects. Sex was reported for a subset of samples within this population (103 females, 227 males).

### ABCC4 Immunoblot

HBEC-6KT cells (n=3), Calu-3 cells (n=3), and primary human bronchial epithelial cells (n=2) were grown to confluence and lysed using RIPA Lysis Buffer containing Protease Inhibitor Cocktail powder for 60-90 min at 4°C on a rocker. Lysates were then centrifuged at 16,000xg for 15 min, and the supernatants were collected. Protein quantification was performed using a BCA protein assay. Equal masses (20μg) of protein were incubated in 1X Laemmli buffer with 0.1 M dithiothreitol at 65°C for 15 min and electrophoresed on 4-15% gradient TGX gels and transferred to PVDF membranes (Bio-Rad Laboratories, Mississauga, Canada). The membranes were blocked with Tris-buffered saline with 0.05% Tween 20 (TBST), and 5% skim milk powder for 2 hours at 25°C. The membranes were incubated with ABCC4/MRP4 (1:40 - M41-10 - Ab15602)). The membranes were washed in TBST, then incubated with horseradish peroxidase-linked anti-rat secondary antibody (1:3000, Cell Signaling Technology^®^, 7077S) or anti-mouse secondary antibody (1:3000, Cell Signaling Technology^®^, 7076S) for 2 hours at 25°C. A chemiluminescence image of the ABCC4 blot was taken in tandem with an image of total protein loading using Bio-Rad Image Lab software (Bio-Rad Laboratories, Mississauga, Canada).

### *In Vitro* Experiments

All experiments were performed with 85-95% confluent HBEC-6KT cells. Cells were treated with interventions (LABA/GCS, ABCC4 inhibitor, and/or PDE4 inhibitors) and incubated at 37°C for 2 hours. After incubation, Poly I:C or imiquimod was added to the cells as a viral mimic challenge. Cell supernatants were collected 24-hour post stimulation and used for quantification of IL-6, IL-8, CCL5/RANTES, and CXCL10/IP-10 using commercially available ELISA kits (R&D Systems, Minnesota, USA). In select experiments, an ABCC4 inhibitor (CF1[10μM]), or PDE4 inhibitors (rolipram[10μM], roflumilast[1μM], and cilomilast[1μM]) were added to LABA/GCS alone or in combination (Ceefourin-1 and roflumilast) followed by viral mimic challenge.

### Statistical Analysis

For the GEO datasets, gene expression levels for ABCC4 were tested for significant differences via Student’s T-test with a Benjamini-Hochberg multiple testing correction using the stats R package (version 3.6.1; R Core Team, 2019). CDH1 and CD34 were used as high and low expression controls, respectively. Gene expression box plots were generated with the ggplot2 R package (version 3.2.1). For the *in vitro* experiments, one-way ANOVAs were performed with a *post-hoc* Bonferroni correction for multiple comparisons and Student’s T-test were performed for single comparisons. Results were represented as means or were normalized as percentages compared to the positive control. A *p*-value <0.05 was accepted to be a statistically significant difference between groups. Data was analyzed using GraphPad Prism Version 8. Data are expressed as mean or mean percentage to control + SEM.

## RESULTS

### ABCC4 expression and localization at gene and protein level

We have demonstrated ABCC4 gene expression in primary human airway epithelial cell cultures grown under submerged monolayer conditions (35). To confirm gene expression in human lung airway epithelium, we performed *in situ* hybridization using RNAscope™ probes on formalin fixed paraffin embedded human lung samples (representative image of n=10). ABCC4 gene transcript was identified as red punctate dots throughout the cytoplasm of airway epithelial cells (**Figure 1A and 1B**). To complement the *in-situ* localization of ABCC4 mRNA transcript in human airway epithelial cells, we mined publicly available gene expression data from bronchial brushings from healthy subjects (n=616). Consistent with our localization studies, ABCC4 gene expression was detected in bronchial brushings at levels between the high control of CDH1 and low control of CD34, genes that were selected to provide reference points for ABCC4 expression levels (**Figure 1C**). No sex dependent gene expression patterns were observed for ABCC4 (**Figure 1D**). To extend our characterization of ABCC4 gene expression, we performed immunohistochemical of ABCC4 protein in human lung tissues (representative image of n=10). Positive staining was observed in human airway epithelial cells (**Figure 1E and F**). *In vitro* confirmation of *in situ* protein expression was performed by immunoblot of cell lines (HBEC-6KT and Calu-3 cells) and primary human airway epithelial cells. Consistent with *in situ* ABCC4 protein expression data, for each airway epithelial cell type, a single band was observed at the predicted molecular weight of 150kDa for ABCC4. Collectively our *in vitro* and *in situ* data confirm gene and protein expression of ABCC4 in human airway epithelial cells.

**Figure 1:**
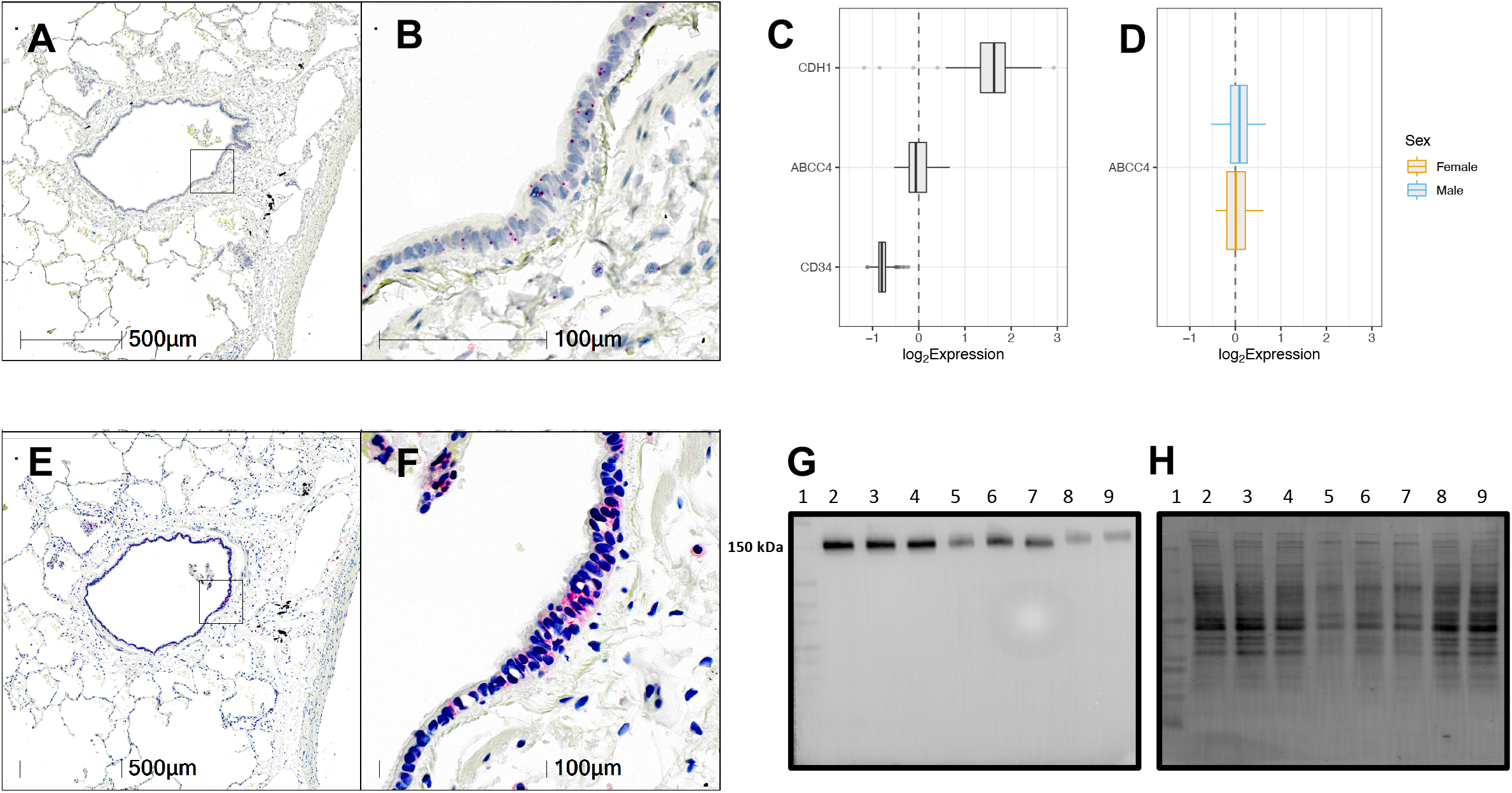
Characterization of ABCC4 expression and localization patterns at the gene and protein level. *In situ* hybridization of ABCC4 RNAscope™ probe in human lung under low (10X) (**A**) and high (40X) (**B**) magnification. Red puncta are representative of ABCC4 gene transcripts with nuclei counterstained blue. Representative image of n=10. (**C**) Gene expression data for 616 healthy subjects with no history of smoking or chronic respiratory disease. (**D**) Healthy samples with metadata defining sex were further divided into male (N=227) and female (N=103) groups and plotted separately as blue and orange-outlined box plots, respectively. For both (C) and (D), log_2_-transformed expression values were plotted as box plots. The dashed line at zero represents the global baseline of expression for the entire set of genes. Immunohistochemistry of ABCC4 in human lung under low (10X) (**E**) and high (40X) (**F**) magnification. Representative image of n=10. Pink/red staining is representative of ABCC4 protein with nuclei counterstained blue. (**G**) ABCC4 protein expression human airway epithelial cells via immunoblot. Lane 1: Protein Ladder, lane 2-4: HBEC-6KT cell lysate (n=3 independent cultures), lane 5-7: Calu-3 cell lysate (n=3 independent cultures), lane 8-9: Primary cell lysate (n=2 independent donors). (H) A total protein stain was performed to demonstrate protein loading levels for the ABCC4 immunoblot.

### Effect of ABCC4 inhibition on LABA/GCS responses in human airway epithelial cells exposed to Poly I:C

We have reported that ABCC4 in human airway epithelial cells functions as a cAMP transporter and may modulate TNF-α-induced responses and efficacy of LABA/GCS interventions (33,35). To interrogate whether ABCC4 played a role in additional innate immune responses in human airway epithelial cells, we performed concentration-response studies with select inflammatory stimuli to define exposure conditions that could be modulated with LABA/GCS. Poly I:C, a TLR3 agonist, induced a concentration-dependent increase in IL-6 and IL-8 (**Figure 2A-B**). In contrast, IL-1β induced an increase in IL-6 and IL-8 at all concentrations examined without a graded response (**Figure 2C-D**). As a result, for all subsequent pharmacological intervention studies, Poly I:C was used as the selected stimulus, at a sub-maximal concentration (1μg/ml) to ensure IL-6 and IL-8 cytokine responses were not saturated and amenable for testing the efficacy of ABCC4 inhibition strategies.

**Figure 2:**
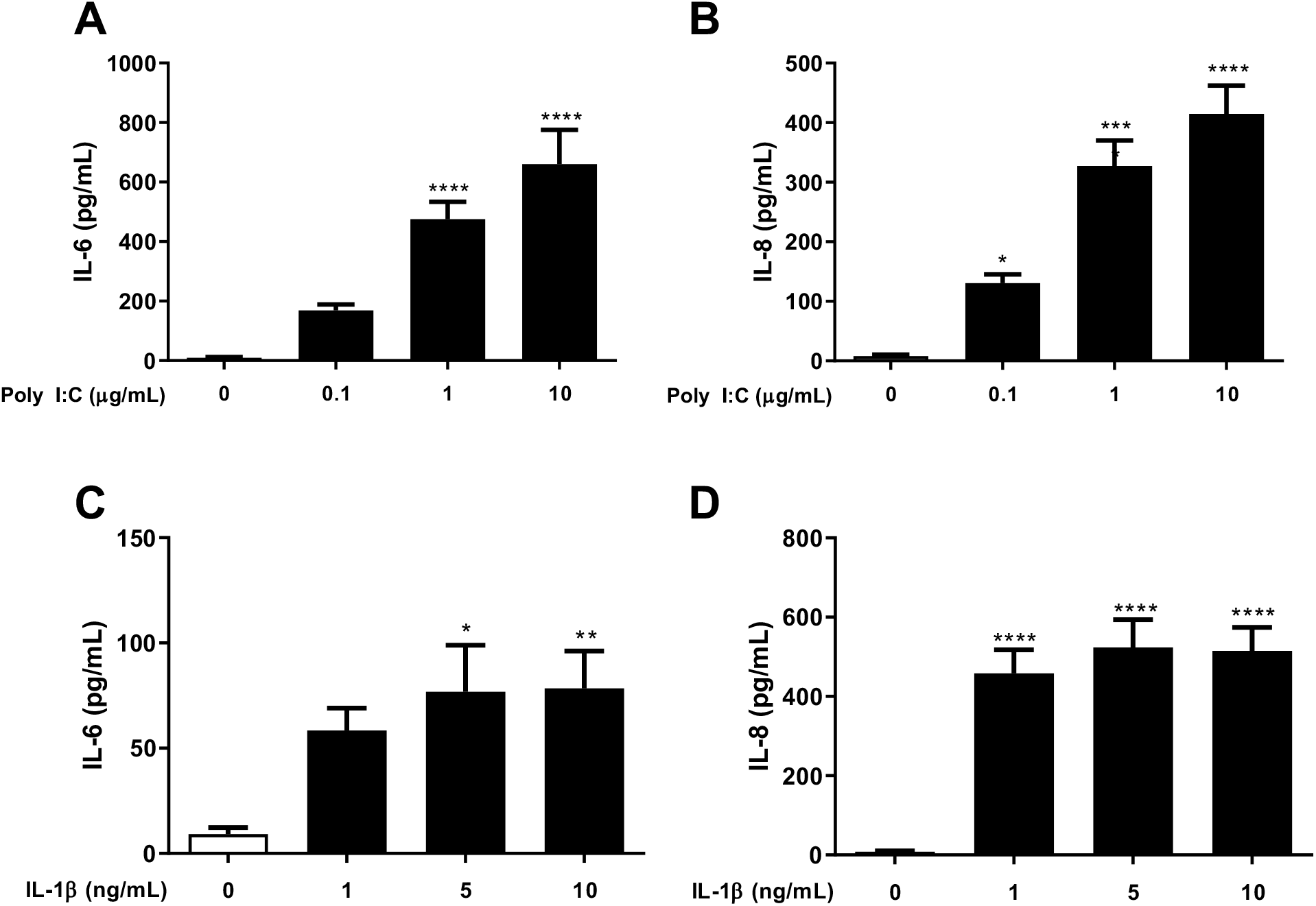
Concentration-response analysis of cytokines response to Poly I:C and IL-1β in human airway epithelial cells. Human airway epithelial cells were exposed to increasing concentrations of Poly I:C (0.1, 1, 10μg/mL), or IL-1β (1, 5, 10 ng/mL), and IL-6, and IL-8 inflammatory cytokines were measured by ELISA. Poly I:C-induced (**A**) IL-6, and (**B**) IL-8. IL-1β-induced (**C**) IL-6, and (**D**) IL-8. Results are shown as mean + SEM. (*p* < 0.05*, *p*<0.01**, *p*<0.001***,*p* < 0.0001**** compared with negative control). n=8.

Our previous report demonstrated that the addition of an ABCC4 inhibitor to LABA/GCS was able to potentiate glucocorticoid response element promoter activity and expression of a LABA/GCS sensitive gene, *RGS2*, in a cAMP-dependent mechanism (35). ABCC4 inhibition potentiates intracellular cAMP levels and contributes to elevated PKA activity (35), which may only be observed on a background of sub-maximal cAMP generation (27). We therefore performed a titration experiment with a fixed concentration of GCS (budesonide – 10nM) and a variable concentration of LABA (formoterol – 0.0001nM to 1nM) under optimized Poly I:C stimulation (1μg/ml). Poly I:C induced both IL-6 and IL-8 responses, which were progressively attenuated with increasing concentrations of formoterol beginning at 0.01nM (**Figure 3A-B**). We therefore determined that 0.01nM formoterol was a sub-maximal concentration to use for ABCC4 intervention studies to ensure that cAMP signaling was not saturated in our human airway epithelial cell model system.

**Figure 3:**
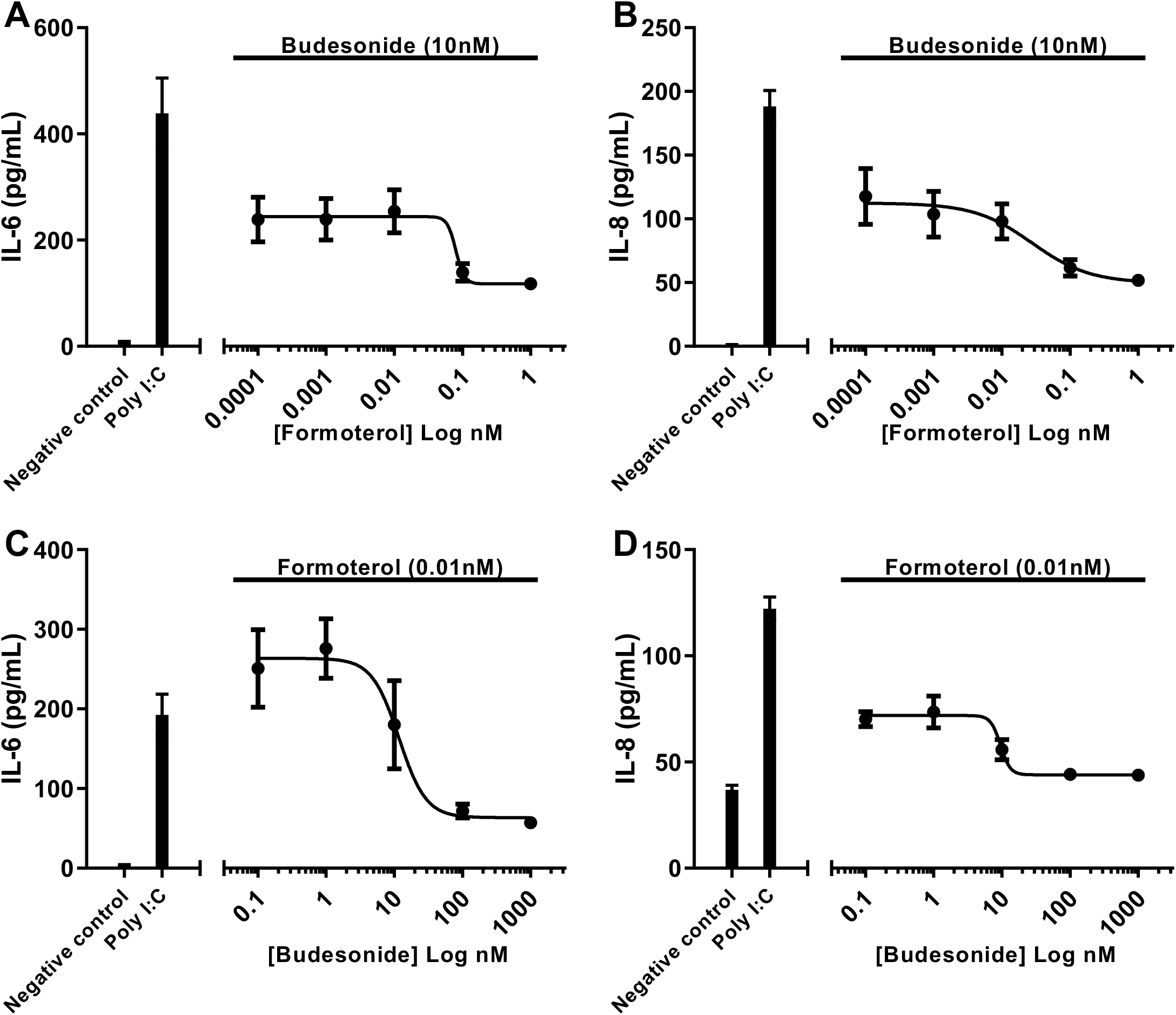
Concentration-response analysis of budesonide and formoterol attenuation of Poly I:C-induced cytokines responses in human airway epithelial cells. Concentration-response analysis measuring (**A**) IL-6 and (**B**) IL-8 was conducted for cells exposed to Poly I:C (1μg/mL) treated with fixed concentration of budesonide (10nM) with increasing log concentrations of formoterol (0.0001, 0.001, 0.01, 0.1, and 1nM). Concentration-response analysis measuring (**C**) IL-6 and (**D**) IL-8 for cells exposed to Poly I:C (1μg/mL) treated with fixed concentration of formoterol (0.01nM) with increasing log concentrations of budesonide (0.1, 1, 10, 100, and 1000nM). Results are shown as mean + SEM. The standard error of means is represented by the error bars attached to each point. n=4.

**Figure 4:**
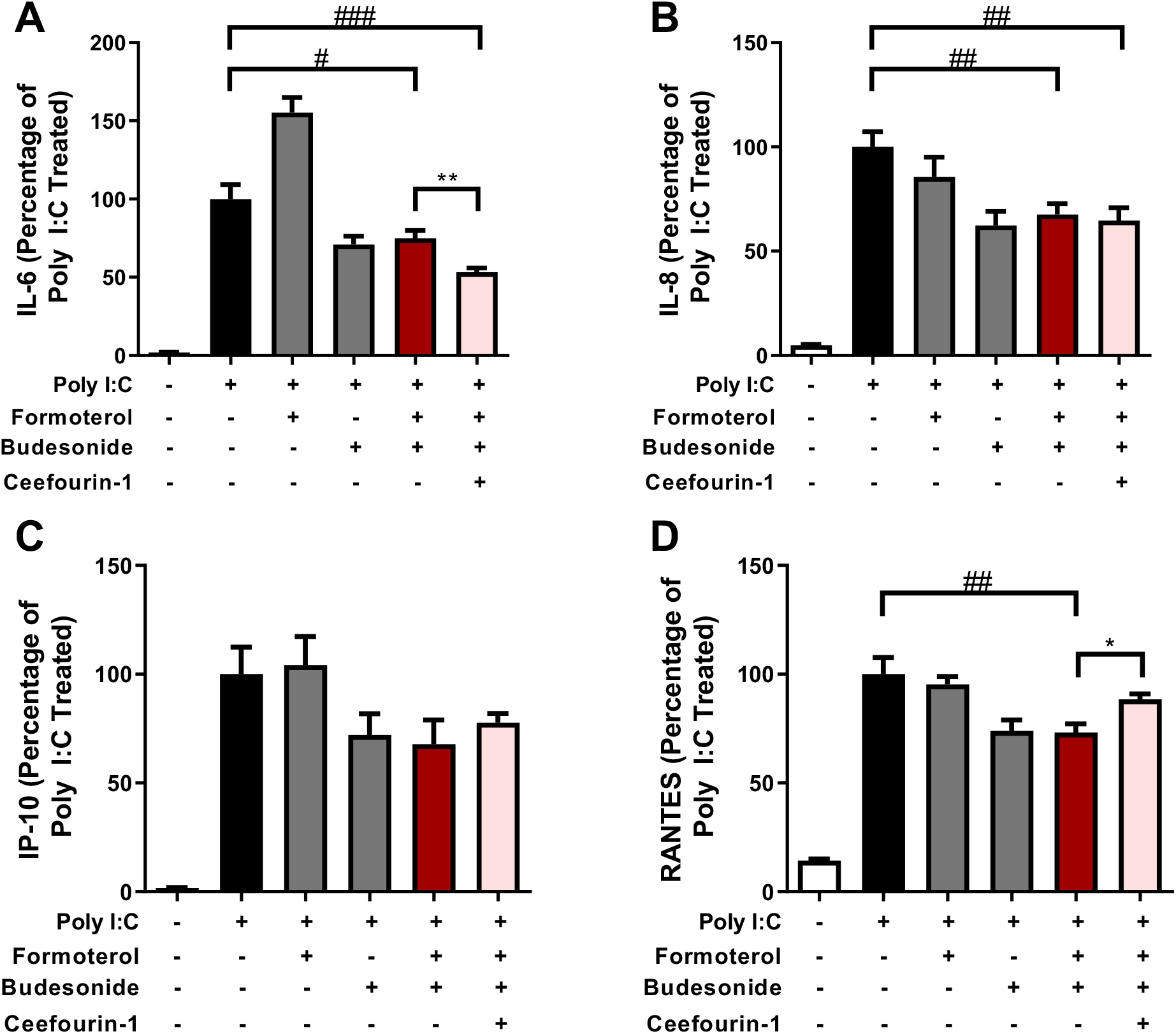
Effect of ABCC4 inhibition on LABA/GCS responses in human airway epithelial cells exposed to Poly I:C. An inflammatory stimulus of Poly I:C (1μg/mL) was used to model a viral challenge in human airway epithelial cells. Combinations of LABA/GCS (budesonide [10nM]/formoterol [0.01nM]) and ABCC4 inhibitor (Ceefourin-1 [10uM]) were performed with Poly I:C exposure and the cytokines, (**A**) IL-6, (**B**) IL-8, (**C**) CXCL10/IP-10, and (**D**) CCL5/RANTES were measured. All data are represented as percentages normalized to positive control stimuli + SEM. (*p* <0.05#, *p* <0.01##, *p* <0.001### compared with Poly I:C, and *p* < 0.05*, *p* <0.01** compared with Poly I:C+budesonide/formoterol). n=5.

Having established a sub-maximal LABA (formoterol) concentration for attenuating Poly I:C-induced IL-6 and IL-8 cytokine responses, we next determined the sub-maximal concentration of GCS (budesonide). Using a similar titration experiment approach, we fixed the formoterol concentration at 0.01nM and used a variable concentration of budesonide (0.1nM to 1000nM). Similar to LABA experiments, Poly I:C induced both IL-6 and IL-8 responses, which were progressively attenuated with increasing concentrations of budesonide beginning at 10nM (**Figure 3C-D**). We therefore determined that 10nM budesonide was a sub-maximal concentration to use for ABCC4 intervention studies to ensure that GCS signaling was not saturated in our human airway epithelial cell model system.

We next combined our optimized experimental conditions of Poly I:C-induced cytokine responses with sub-maximal formoterol and budesonide concentrations, with a selective ABCC4 intervention. To inhibit ABCC4, we used the selective ABCC4 small molecule inhibitor, Ceefourin-1, at 10uM, to approximate an IC_50_ concentration reported in HEK cells (37). Budesonide and formoterol were fixed at 10nM and 0.01nM, respectively. LABA/GCS suppressed Poly I:C-induced IL-6 by 25.0% compared to the positive control and addition of Ceefourin-1 was able to suppress Poly I:C-induced IL-6 by 46.8% compared to positive control (*p* <0.05 compared to LABA/GCS) (**Figure 5A**). The addition of Ceefourin-1 did not significantly impact IL-8 (**Figure 5B**). As Poly I:C is a TLR3 agonist and viral mimic, we also interrogated CXCL10/IP-10 and CCL5/RANTES to determine if additional cytokines important in antiviral responses were sensitive to ABCC4 inhibition in combination with LABA/GCS. Similar to IL-8, ABCC4 inhibition did not potentiate LABA/GCS attenuation of Poly I:C-induced CXCL10/IP-10 or CCL5/RANTES (**Figure 5C-D**).

**Figure 5.**
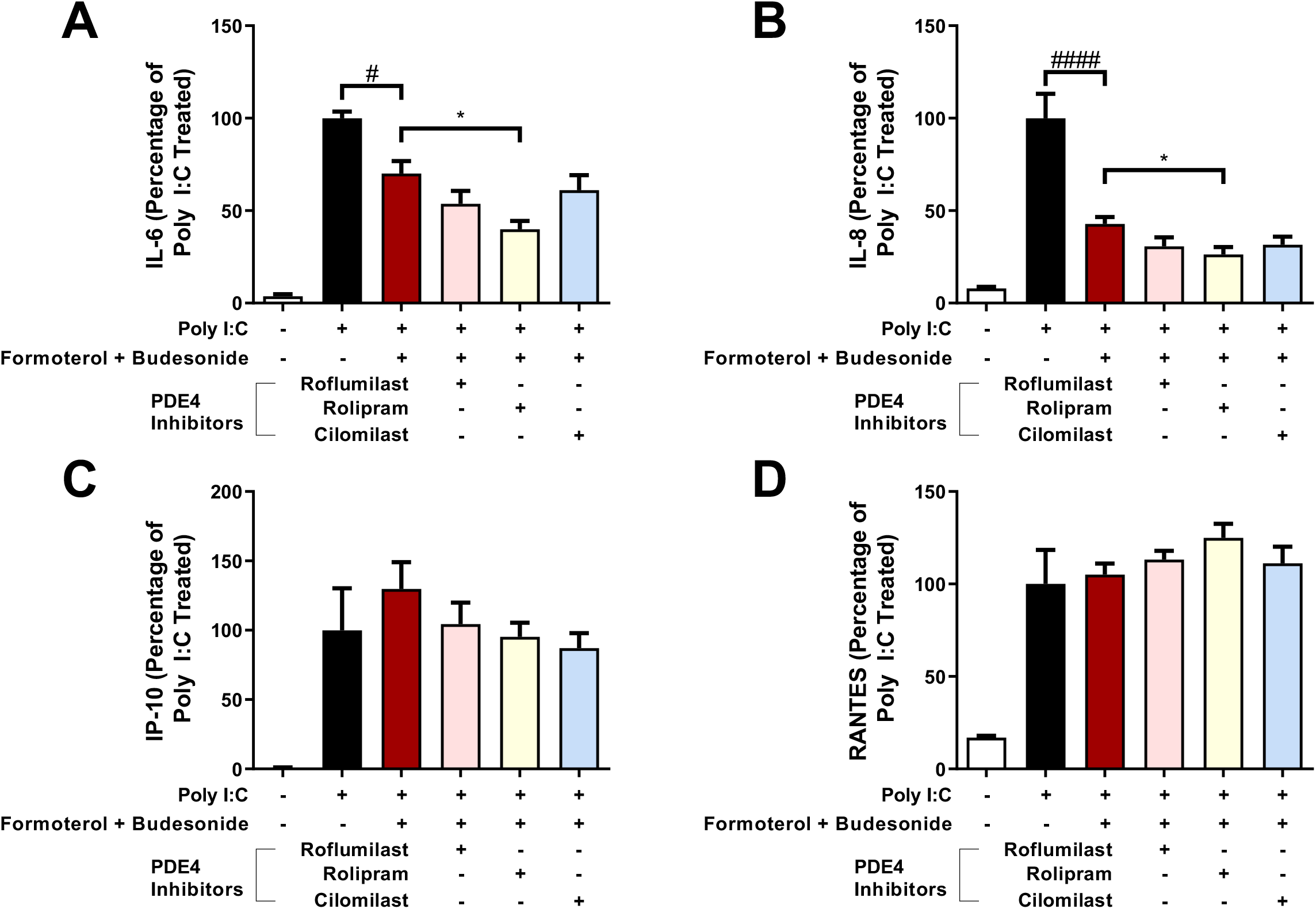
Effect of PDE4 inhibition on LABA/GCS responses in human airway epithelial cells exposed to Poly I:C. An inflammatory stimulus of Poly I:C (1μg/mL) was used to model a viral challenge in human airway epithelial cells. Combinations of LABA/GCS (budesonide [10nM]/formoterol [0.01nM]) and PDE4 inhibitors (roflumilast [1uM], rolipram [10uM], cilomilast [1uM]) were performed with Poly I:C exposure and the cytokines, (**A**) IL-6, (**B**) IL-8, (**C**) CXCL10/IP-10, and (**D**) CCL5/RANTES were measured. All data are represented as percentages normalized to positive control stimuli + SEM. (*p* <0.05#, *p* <0.0001#### compared with Poly I:C, and *p* < 0.05* compared with Poly I:C+budesonide/formoterol) n=5.

### Effect of PDE4 inhibition on LABA/GCS responses in human airway epithelial cells exposed to Poly I:C

PDE4 inhibition represents a clinically approved strategy for elevating intracellular cAMP for management of chronic respiratory diseases. To complement and provide context for our ABCC4 inhibition studies we performed combination LABA/GCS and PDE4 inhibition experiments. Using the same method as ABCC4 inhibition experiments, we substituted the ABCC4 inhibitor with three distinct PDE4 inhibitors, roflumilast, rolipram, and cilomilast. LABA/GCS suppressed Poly I:C-induced IL-6 by 29.9% compared to the positive control. Addition of rolipram was able to suppress Poly I:C-induced IL-6 by 60.1% compared to positive control (*p*<0.05 compared to LABA/GCS), but the addition of roflumilast or cilomilast did not significantly impact IL-6 levels (**Figure 6A**). LABA/GCS suppressed Poly I:C-induced IL-8 by 57.2% compared to the positive control. Addition of rolipram was able to suppress Poly I:C-induced IL-8 by 73.7% compared to positive control (*p*<0.05 compared to LABA/GCS), but the addition of roflumilast or cilomilast did not significantly impact IL-8 levels (**Figure 6B**). The addition of any PDE4 inhibitors did not significantly attenuate CXCL10/IP-10 or CCL5/RANTES when compared to both the positive control or LABA/GCS (**Figure 6C and Figure 6D**).

**Figure 6.**
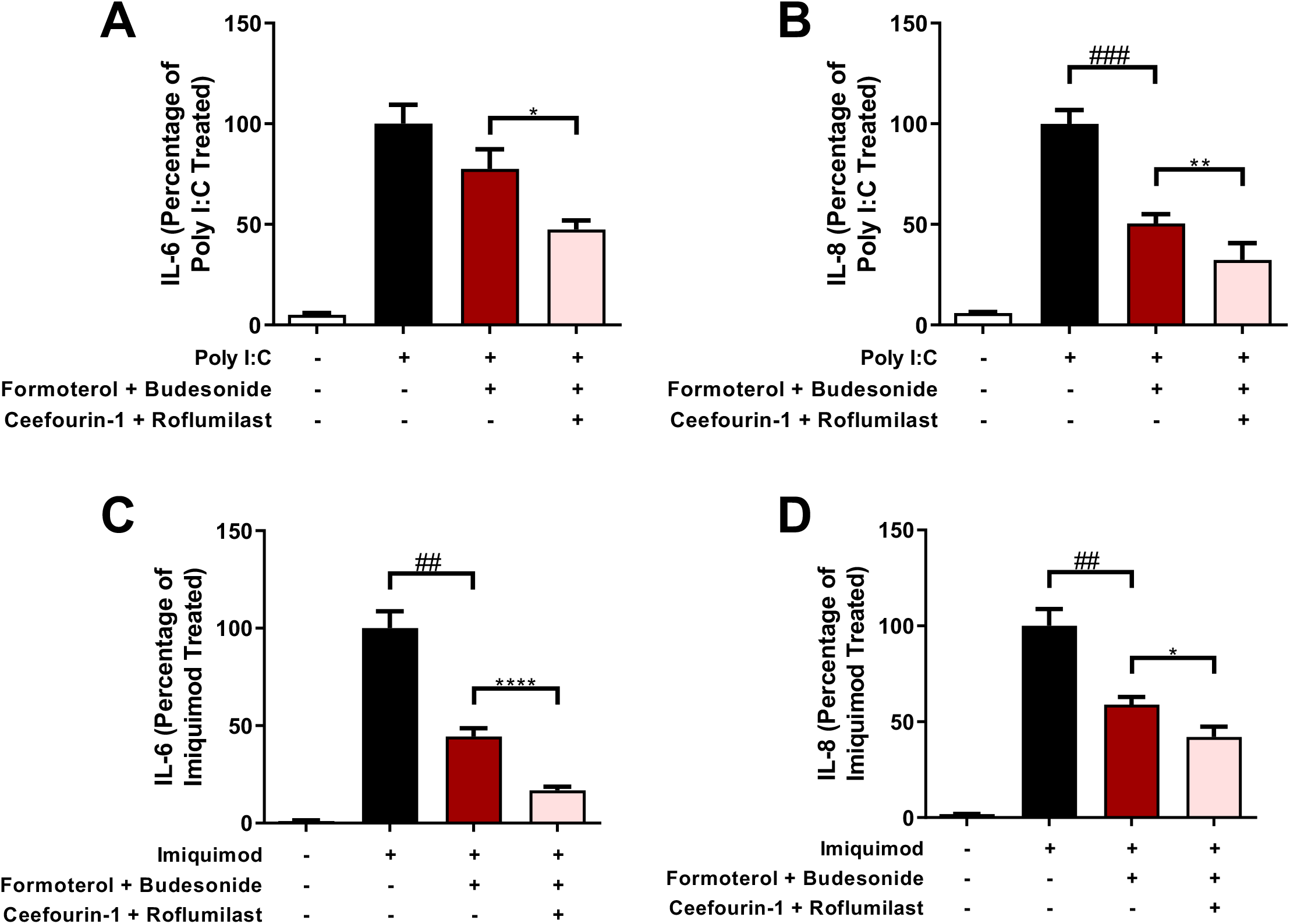
Effect of combination ABCC4 and PDE4 inhibition on LABA/GCS responses in human airway epithelial cells exposed to TLR ligands. An inflammatory stimulus of Poly I:C (1μg/mL) was used to model a viral challenge in human airway epithelial cells. Combinations of LABA/GCS (budesonide [10nM]/formoterol [0.01nM]), ABCC4 inhibitor (Ceefourin-1[10uM]) and PDE4 inhibitor (roflumilast [1uM]) were performed with Poly I:C exposure and the cytokines, (**A**) IL-6, (**B**) IL-8 were measured (n=6). Then, using the same conditions but changing the inflammatory stimulus to imiquimod (100μg/mL) (**C**) IL-6, (**D**) IL-8 were measured (n=4). All data are represented as percentages normalized to Poly I:C/imiquimod + SEM. (*p* <0.01##, *p* <0.001### compared with positive control, and *p* < 0.05*, *p* <0.01**, *p* <0.0001**** compared with Poly I:C/imiquimod+budesonide/formoterol).

### Effect of combination ABCC4 and PDE4 inhibition on LABA/GCS responses in human airway epithelial cells exposed to TLR ligands

Regulation of intracellular cAMP levels may be co-regulated by both extracellular transport (i.e. ABCC4) and enzymatic degradation (i.e. PDE4) (27). Inhibition of only one arm of the intracellular cAMP regulation processes may shunt cAMP to the other arm. We therefore performed combination ABCC4 and PDE4 inhibition experiments with LABA/GCS in the presence of Poly I:C. Using the same methods as above, we compared the difference between LABA/GCS alone and LABA/GCS with both ABCC4 (Ceefourin-1) and PDE4 inhibition, using roflumilast due to clinical approval and use. LABA/GCS suppressed Poly I:C-induced IL-6 by 22.5%, compared to the positive control (**Figure 7A**). Addition of Ceefourin-1+roflumilast was able to suppress Poly I:C-induced IL-6 by 52.5% compared to positive control (*p*<0.05 compared to LABA/GCS). LABA/GCS suppressed Poly I:C-induced IL-8 by 49.5%, compared to the positive control (**Figure 7B**). Addition of Ceefourin-1+roflumilast was able to suppress Poly I:C-induced IL-8 by 67.7% compared to positive control (*p*<0.01 compared to LABA/GCS). Similar to ABCC4 and PDE4 experiments, CXCL10/IP-10 or CCL5/RANTES were not impacted by combination intervention (data not shown).

Poly I:C is a TLR3 ligand used as a viral mimic (50). To determine whether our observed cAMP modulation results were conserved across different TLR pathways related to viral stimulation, we subsequently performed intervention experiments with imiquimod, a TLR7 ligand, in place of Poly I:C. LABA/GCS suppressed imiquimod-induced IL-6 by 55.6%, compared to the positive control, and addition of Ceefourin-1+roflumilast was able to suppress imiquimod-induced IL-6 by 83.3% compared to positive control (*p*<0.001 compared to LABA/GCS) (**Figure 7C**). Imiquimod-induced IL-8 was also suppressed by LABA/GCS (41.1%) compared to the positive control, and addition of Ceefourin-1+roflumilast further suppressed IL-8 (57.9%) compared to positive control (*p*<0.01 compared to LABA/GCS) (**Figure 7D**).

## DISCUSSION

Our study investigated the impact of cAMP modulation via inhibiting extracellular transport or intracellular breakdown on LABA/GCS efficacy in human airway epithelial cells using a combinatorial pharmacological approach. We confirmed the expression of ABCC4 in human airway epithelial cells and its function in the context of regulating immune responses relevant in asthma and chronic respiratory disease (33,35). We also demonstrated ABCC4 or PDE4 inhibition potentiated LABA/GCS attenuation of Poly I:C-induced IL-6 and IL-8, which was greater when ABCC4 and PDE4 inhibition was combined. Modulation of cAMP levels by all combinatorial approaches had no impact on LABA/GCS modulation of Poly I:C-induced CXCL10/IP-10 or CCL5/RANTES. The observed potentiation of LABA/GCS efficacy by ABCC4 and PDE4 combination was conserved beyond a TLR3/Poly I:C pathway, with similar observations made with TLR7/imiquimod. Importantly, we also independently confirm previous reports of formoterol potentiation of IL-6 responses in human airway epithelial cells during exposure to viral stimuli (17,18), further cautioning against using cAMP-elevating agents in isolation.

The modulation of intracellular cAMP levels can be achieved by activation of adenylyl cyclases, inhibition of PDEs, or preventing extracellular transport by ABCC4 (27). Increasing intracellular cAMP levels contributes to elevated PKA activity and may synergize with GCS anti-inflammatory responses (36). We have previously demonstrated that inhibition of ABCC4 with the non-selective inhibitor, MK-571, was able to potentiate PKA activity and LABA/GCS responses (35). However, in this study we utilized Ceefourin-1, a selective ABCC4 inhibitor that does not include the PDE4 inhibitor properties of MK-571 (37). Using Ceefourin-1 at IC_50_ concentrations, we observed a significant potentiation of LABA/GCS attenuation of Poly I:C-induced IL-6, but not IL-8. The results observed with Ceefourin-1 were similar in direction as with all three PDE4 inhibitors examined, suggesting that either mechanism of inhibition of extracellular transport or intracellular metabolism, could contribute to LABA/GCS potentiation.

PDE4 and ABCC4 have different subcellular localization. PDE4 is present throughout the cell with different isoforms localized to different cellar compartments such as the cytosol, Golgi body, and nucleus as well as being coupled to plasma membrane proteins (51,52). In contrast, ABCC4 is confined to the plasma membrane, which could indicate the presence of a more compartmentalized increase in cAMP. A study looking at the spatiotemporal coupling of ABCC4 to CFTR in gut epithelial cells have shown that blocking ABCC4 enhances CFTR function at lower concentrations of adenosine, an adenylyl cyclase agonist, but not at higher concentrations (53). This data supports a model in which blocking ABCC4 leads to compartmentalized cAMP accumulation rather then a global cellular increase in cAMP. A model that is further buttressed by the fact that overexpression of ABCC4 does not lead to a substantial decrease in intracellular cAMP levels (54). As ABCC4 and PDE4 may function within different cellular compartments (27), our data suggest that global cAMP levels, and not cell compartment specific, are able to potentiate LABA/GCS regulation of IL-6 and IL-8.

To expand our analysis beyond pro-inflammatory cytokines, we considered cytokines involved with antiviral responses, analyzing CXCL10/IP-10 and CCL5/RANTES in experiments with ABCC4 and PDE4 inhibitors. We observed LABA/GCS attenuation of Poly I:C-induced IP-10 and RANTES levels, which tended to increase with ABCC4 inhibition. Consistent with ABCC4 inhibition, PDE4 inhibition also revealed a trend to potentiate RANTES, but not IP-10. These data demonstrate that unlike IL-6 or IL-8, which were attenuated to a greater extent during LABA/GCS treatment combined with mechanisms that further increase intracellular cAMP, this may not be a global response for all cytokines. Indeed, this is consistent with transcriptomic responses in human airway epithelial cells exposed to LABA/GCS, where some GCS regulated genes are variably enhanced, repressed, or unaffected by the co-administration of LABA (55). The differential direction for regulation of cytokine responses by LABA/GCS combinatorial treatment with ABCC4 or PDE4 inhibitors is important as suppression of a particular pathway examined may concomitantly be accompanied by potentiation of another. Our results suggest that suppression of pro-inflammatory cytokine responses via cAMP modulation and GCS could be accompanied with potentiation of antiviral cytokine levels. Although the expression of DUSP1 is able to repress the expression of many inflammatory genes, it has been shown that IL-1β-induced IP-10 gene expression is further increased upon DUSP1overexpression. A potential reason for this could be because DUSP1 maintains expression of interferon regulatory factor 1 (IRF-1) and IRF1-dependent gene, IP-10 (56). RANTES has been shown to be expressed at an elevated level in asthmatic children compared to healthy children, and in studies conducted in adult asthmatics shows increased RANTES during asthma exacerbations (57–60). Our observations that IP-10 and RANTES are not significantly attenuated with cAMP modulation (either ABCC4 or PDE4 interventions) on a background of LABA/GCS treatment suggest that a host antiviral response mediated by these antiviral cytokines is maintained.

PDE4 inhibitors have been explored and approved (e.g. roflumilast) for clinical use of chronic respiratory disease, operating by a mechanism of elevating intracellular cAMP levels. PDE4 inhibitors are frequently associated with a high incidence of side effects such as nausea and vomiting (61) which may be related to the broad distribution of PDE4 enzymes throughout the GI tract, an observation remaining to be determined for ABCC4. Based on previous literature that evaluated their anti-inflammatory effects in human airway epithelial cells, we chose 1μM for roflumilast, 10μM for rolipram, and 1μM for cilomilast (26,62,63). Roflumilast, the clinically available drug, was not able to significantly alter LABA/GCS effectiveness in any of the cytokine measurements. This trend was also seen in cilomilast. However, rolipram was able to further suppress both IL-6 and IL-8 levels. CXCL10/IP-10 and CCL5/RANTES readout were not further augmented with the addition of any of the PDE4 inhibitors. The difference in the PDE4 inhibitors may be due to its preferential selectivity to specific isoforms. Rolipram favourably inhibits PDE4A (IC_50_ = 3nM) over other isoforms such as B and D (IC_50_ = 130nM and IC_50_ = 240nM respectively) (64). Whereas, roflumilast has a preferential inhibition of PDE4B and PDE4D (IC_50_ < 1nM for both) (65). To support this claim, cilomilast also shows 10-fold selectivity toward PDE4D compared with PDE4A and PDE4B (66). Although clinically, PDE4 inhibitors are efficacious, there are side effects that are frequently associated with them. However, there is a possibility that combining or replacing PDE4 inhibitors with ABCC4 inhibitors may help minimize the side-effects by lowering doses or targeting cAMP modulation away from the GI tract (67).

Inhibiting ABCC4 and PDE4 independently may lead to shunting of cAMP to the alternate metabolic pathway (27). To account for this possibility, we inhibited ABCC4 extracellular transport of cAMP in combination with inhibition of cAMP metabolism by PDE4. For the PDE4 inhibitor, we selected roflumilast since it is approved for clinical use. We performed these experiments with both Poly I:C and imiquimod to interrogate whether different TLR agonist-induced cytokine responses were equally impacted by modulating cAMP levels. Poly I:C and imiquimod as stimuli, both IL-6 and IL-8 were further suppressed whereas in Poly I:C. The data demonstrate that ABCC4 and PDE4 inhibitors work together to increase intracellular cAMP since Ceefourin-1 and roflumilast by themselves were not able to augment IL-8 levels, but in combination were able to significantly further decrease IL-8 compared to LABA/GCS.

In conclusion, we demonstrate that modulation of intracellular cAMP levels in human airway epithelial cells with ABCC4 and PDE4 interventions could potentiate the anti-inflammatory effects of LABA/GCS for Poly I:C and imiquimod-induced inflammation. Our findings complement current literature which shows that ABCC4 inhibition potentiates LABA/GCS by the up-regulation of anti-inflammatory and bronchoprotective genes. Our results form the basis to further explore the potential application of ABCC4 and PDE4 inhibitors for attenuation of inflammation in the context of chronic respiratory diseases.

## Notes

### Competing Interest Statement

The authors have declared no competing interest.

## References

1. Roth GA, Abate D, Abate KH, Abay SM, Abbafati C, Abbasi N, et al. Global, regional, and national age-sex-specific mortality for 282 causes of death in 195 countries and territories, 1980-2017: a systematic analysis for the Global Burden of Disease Study 2017. Lancet. 2018;

2. Martin RJ, Szefler SJ, King TS, Kraft M, Boushey HA, Chinchilli VM, et al. The Predicting Response to Inhaled Corticosteroid Efficacy (PRICE) trial. J Allergy Clin Immunol. 2007;

3. Szefler SJ, Martin RJ, King TS, Boushey HA, Cherniack RM, Chinchilli VM, et al. Significant variability in response to inhaled corticosteroids for persistent asthma. J Allergy Clin Immunol. 2002;109(3):410–8.

4. Loftus PA, Wise SK. Epidemiology of asthma. Current Opinion in Otolaryngology and Head and Neck Surgery. 2016.

5. Public Health Agency of Canada. Report from the Canadian Chronic Disease Surveillance System: Asthma and Chronic Obstructive Pulmonary Disease (COPD) in Canada. [Internet]. 2018. Available from: https://www.canada.ca/content/dam/phac-aspc/documents/services/publications/diseases-conditions/asthma-chronic-obstructive-pulmonary-disease-canada-2018/pub-eng.pdf

6. Global Initiative for Asthma. Pocket Guide for Asthma Management and Prevention. Global Initiative for Asthma. 2019.

7. Wu P, Hartert T V. Evidence for a causal relationship between respiratory syncytial virus infection and asthma. Expert Review of Anti-Infective Therapy. 2011.

8. Dahlén B, Roquet A, Inman MD, Karlsson Ö, Naya I, Anstrén G, et al. Influence of zafirlukast and loratadine on exercise-induced bronchoconstriction. J Allergy Clin Immunol. 2002;

9. O’byrne PM, Inman MD, Parameswaran K. The trials and tribulations of IL-5, eosinophils, and allergic asthma. J Allergy Clin Immunol. 2001;

10. Johnston NW, Johnston SL, Duncan JM, Greene JM, Kebadze T, Keith PK, et al. The September epidemic of asthma exacerbations in children: A search for etiology. J Allergy Clin Immunol. 2005;

11. Jackson DJ, Sykes A, Mallia P, Johnston SL. Asthma exacerbations: Origin, effect, and prevention. J Allergy Clin Immunol. 2011;

12. Papadopoulos NG, Papi A, Psarras S, Johnston SL. Mechanisms of rhinovirus-induced asthma. Paediatr Respir Rev. 2004;

13. Barnes PJ. How corticosteroids control inflammation: Quintiles Prize Lecture 2005. British Journal of Pharmacology. 2006.

14. Newton R, Holden NS. Separating transrepression and transactivation: A distressing divorce for the glucocorticoid receptor? Molecular Pharmacology. 2007.

15. Dirks NL, Li S, Huth B, Hochhaus G, Yates CR, Meibohm B. Transrepression and transactivation potencies of inhaled glucocorticoids. Pharmazie. 2008;

16. Johnson M. Molecular mechanisms of β2-adrenergic receptor function, response, and regulation. Journal of Allergy and Clinical Immunology. 2006.

17. Ritchie AI, Singanayagam A, Wiater E, Edwards MR, Montminy M, Johnston SL. Beta2-Agonists Enhance Asthma-Relevant Inflammatory Mediators in Human Airway Epithelial Cells. American Journal of Respiratory Cell and Molecular Biology. 2018.

18. Edwards MR, Haas J, Panettieri RA, Johnson M, Johnston SL. Corticosteroids and β 2 agonists differentially regulate rhinovirus-induced interleukin-6 via distinct cis-acting elements. J Biol Chem. 2007;

19. Newton R, Giembycz MA. Understanding how long-acting β 2-adrenoceptor agonists enhance the clinical efficacy of inhaled corticosteroids in asthma - an update. British Journal of Pharmacology. 2016.

20. Johnasson-Haque K, Palanichamy E, Okret S. Stimulation of MAPK-phosphatase 1 gene expression by glucocorticoids occurs through a tethering mechanism involving C/EBP. J Mol Endocrinol. 2008;

21. Shipp LE, Lee J V., Yu CY, Pufall M, Zhang P, Scott DK, et al. Transcriptional regulation of human dual specificity protein phosphatase 1 (DUSP1) gene by glucocorticoids. PLoS One. 2010;

22. Tchen CR, Martins JRS, Paktiawal N, Perelli R, Saklatvala J, Clark AR. Glucocorticoid regulation of mouse and human dual specificity phosphatase 1 (DUSP1) genes: Unusual cis-acting elements and unexpected evolutionary divergence. J Biol Chem. 2010;

23. Zhang W, Liu HT. MAPK signal pathways in the regulation of cell proliferation in mammalian cells. Cell Res. 2006;

24. Holden NS, George T, Rider CF, Chandrasekhar A, Shah S, Kaur M, et al. Induction of Regulator of G-Protein Signaling 2 Expression by Long-Acting 2-Adrenoceptor Agonists and Glucocorticoids in Human Airway Epithelial Cells. J Pharmacol Exp Ther [Internet]. 2013;348(1):12–24. Available from: http://jpet.aspetjournals.org/cgi/doi/10.1124/jpet.113.204586

25. Holden NS, Bell MJ, Rider CF, King EM, Gaunt DD, Leigh R, et al. 2-Adrenoceptor agonist-induced RGS2 expression is a genomic mechanism of bronchoprotection that is enhanced by glucocorticoids. Proc Natl Acad Sci. 2011;

26. Moodley T, Wilson SM, Joshi T, Rider CF, Sharma P, Yan D, et al. Phosphodiesterase 4 Inhibitors Augment the Ability of Formoterol to Enhance Glucocorticoid-Dependent Gene Transcription in Human Airway Epithelial Cells: A Novel Mechanism for the Clinical Efficacy of Roflumilast in Severe Chronic Obstructive Pulmonary Di. Mol Pharmacol. 2013;

27. Cheepala S, Hulot J-S, Morgan JA, Sassi Y, Zhang W, Naren AP, et al. Cyclic Nucleotide Compartmentalization: Contributions of Phosphodiesterases and ATP-Binding Cassette Transporters. Annu Rev Pharmacol Toxicol. 2011;

28. Conti M, Beavo J. Biochemistry and Physiology of Cyclic Nucleotide Phosphodiesterases: Essential Components in Cyclic Nucleotide Signaling. Annu Rev Biochem. 2007;

29. Page CP, Spina D. Phosphodiesterase inhibitors in the treatment of inflammatory diseases. Handb Exp Pharmacol. 2011;

30. Halpin DMG. ABCD of the phosphodiesterase family: Interaction and differential activity in COPD. International Journal of COPD. 2008.

31. Russel FGM, Koenderink JB, Masereeuw R. Multidrug resistance protein 4 (MRP4/ABCC4): a versatile efflux transporter for drugs and signalling molecules. Vol. 29, Trends in Pharmacological Sciences. 2008. p. 200–7.

32. Conner GE, Ivonnet P, Gelin M, Whitney P, Salathe M. H2O2 stimulates cystic fibrosis transmembrane conductance regulator through an autocrine prostaglandin pathway, using multidrug-resistant protein-4. Am J Respir Cell Mol Biol. 2013;

33. Gold MJ, Hiebert PR, Park HY, Stefanowicz D, Le A, Starkey MR, et al. Mucosal production of uric acid by airway epithelial cells contributes to particulate matter-induced allergic sensitization. Mucosal Immunol. 2016;9(3):809–20.

34. Ahmadi S, Bozoky Z, Di Paola M, Xia S, Li C, Wong AP, et al. Phenotypic profiling of CFTR modulators in patient-derived respiratory epithelia. npj Genomic Med. 2017;

35. Huff RD, Rider CF, Yan D, Newton R, Giembycz MA, Carlsten C, et al. Inhibition of ABCC4 potentiates combination beta agonist and glucocorticoid responses in human airway epithelial cells. Journal of Allergy and Clinical Immunology. 2017;

36. Giembycz MA, Newton R. Potential mechanisms to explain how LABAs and PDE4 inhibitors enhance the clinical efficacy of glucocorticoids in inflammatory lung diseases. F1000Prime Rep [Internet]. 2015;7. Available from: http://www.f1000.com/prime/reports/b/7/16

37. Cheung L, Flemming CL, Watt F, Masada N, Yu DMT, Huynh T, et al. High-throughput screening identifies Ceefourin 1 and Ceefourin 2 as highly selective inhibitors of multidrug resistance protein 4 (MRP4). Biochem Pharmacol. 2014;

38. Ramirez RD, Sheridan S, Girard L, Sato M, Kim Y, Pollack J, et al. Immortalization of human bronchial epithelial cells in the absence of viral oncoproteins. Cancer Res. 2004;

39. Hirota JA, Gold MJ, Hiebert PR, Parkinson LG, Wee T, Smith D, et al. The nucleotide-binding domain, leucine-rich repeat protein 3 inflammasome/IL-1 receptor I axis mediates innate, but not adaptive, immune responses after exposure to particulate matter under 10 μm. Am J Respir Cell Mol Biol. 2015;

40. Huff RD, Hsu ACY, Nichol KS, Jones B, Knight DA, Wark PAB, et al. Regulation of xanthine dehydrogensase gene expression and uric acid production in human airway epithelial cells. PLoS One. 2017;

41. Hirota JA, Marchant DJ, Singhera GK, Moheimani F, Dorscheid DR, Carlsten C, et al. Urban particulate matter increases human airway epithelial cell IL-1β secretion following scratch wounding and H1N1 influenza A exposure in vitro. Exp Lung Res. 2015;

42. Aguiar JA, Huff RD, Tse W, Stämpfli MR, McConkey BJ, Doxey AC, et al. Transcriptomic and barrier responses of human airway epithelial cells exposed to cannabis smoke. Physiol Rep. 2019;

43. Sean D, Meltzer PS. GEOquery: A bridge between the Gene Expression Omnibus (GEO) and BioConductor. Bioinformatics. 2007;

44. Dai M, Wang P, Boyd AD, Kostov G, Athey B, Jones EG, et al. Evolving gene/transcript definitions significantly alter the interpretation of GeneChip data. Nucleic Acids Res. 2005;

45. Piccolo SR, Sun Y, Campbell JD, Lenburg ME, Bild AH, Johnson WE. A single-sample microarray normalization method to facilitate personalized-medicine workflows. Genomics. 2012;

46. Huber W, Carey VJ, Gentleman R, Anders S, Carlson M, Carvalho BS, et al. Orchestrating high-throughput genomic analysis with Bioconductor. Nat Methods. 2015;

47. Johnson WE, Li C, Rabinovic A. Adjusting batch effects in microarray expression data using empirical Bayes methods. Biostatistics. 2007;

48. Leek JT, Johnson WE, Parker HS, Jaffe AE, Storey JD. The SVA package for removing batch effects and other unwanted variation in high-throughput experiments. Bioinformatics. 2012;

49. Cheadle C, Vawter MP, Freed WJ, Becker KG. Analysis of microarray data using Z score transformation. J Mol Diagnostics. 2003;

50. Lever AR, Park H, Mulhern TJ, Jackson GR, Comolli JC, Borenstein JT, et al. Comprehensive evaluation of poly(I:C) induced inflammatory response in an airway epithelial model. Physiol Rep. 2015;

51. Oldenburger A, Maarsingh H, Schmidt M. Multiple facets of cAMP signalling and physiological impact: CAMP compartmentalization in the lung. Pharmaceuticals. 2012.

52. Conti M, Richter W, Mehats C, Livera G, Park J-Y, Jin C. Cyclic AMP-specific PDE4 Phosphodiesterases as Critical Components of Cyclic AMP Signaling. J Biol Chem. 2003;

53. Li C, Krishnamurthy PC, Penmatsa H, Marrs KL, Wang XQ, Zaccolo M, et al. Spatiotemporal Coupling of cAMP Transporter to CFTR Chloride Channel Function in the Gut Epithelia. Cell. 2007;

54. Wielinga PR, Van der Heijden I, Reid G, Beijnen JH, Wijnholds J, Borst P. Characterization of the MRP4- and MRP5-mediated transport of cyclic nucleotides from intact cells. J Biol Chem. 2003;

55. Rider CF, Altonsy MO, Mostafa MM, Shah S V., Sasse S, Manson ML, et al. Long-acting b 2-Adrenoceptor agonists enhance glucocorticoid receptor (gr)–mediated transcription by gene-specific mechanisms rather than generic effects via GR s. Mol Pharmacol. 2018;

56. Shah S, King EM, Mostafa MM, Altonsy MO, Newton R. DUSP1 maintains IRF1 and leads to increased expression of IRF1-dependent genes: A mechanism promoting glucocorticoid insensitivity. J Biol Chem. 2016;

57. Ying S, Meng Q, Zeibecoglou K, Robinson DS, Macfarlane A, Humbert M, et al. Eosinophil Chemotactic Chemokines (Eotaxin, Eotaxin-2, RANTES, Monocyte Chemoattractant Protein-3 (MCP-3), and MCP-4), and C-C Chemokine Receptor 3 Expression in Bronchial Biopsies from Atopic and Nonatopic (Intrinsic) Asthmatics. J Immunol. 1999;

58. Teran LM, Noso N, Carroll M, Davies DE, Holgate S, Schroder JM. Eosinophil recruitment following allergen challenge is associated with the release of the chemokine RANTES into asthmatic airways. J Immunol. 1996;

59. Lamkhioued B, Renzi PM, Abi-Younes S, Garcia-Zepada EA, Allakhverdi Z, Ghaffar O, et al. Increased expression of eotaxin in bronchoalveolar lavage and airways of asthmatics contributes to the chemotaxis of eosinophils to the site of inflammation. J Immunol. 1997;

60. Lamkhioued B, Garcia-Zepeda EA, Abi-Younes S, Nakamura H, Jedrzkiewicz S, Wagner L, et al. Monocyte chemoattractant protein (MCP)-4 expression in the airways of patients with asthma: Induction in epithelial cells and mononuclear cells by proinflammatory cytokines. Am J Respir Crit Care Med. 2000;

61. Banner KH, Page CP. Theophylline and selective phosphodiesterase inhibitors as anti-inflammatory drugs in the treatment of bronchial asthma. European Respiratory Journal. 1995.

62. BinMahfouz H, Borthakur B, Yan D, George T, Giembycz MA, Newton R. Superiority of Combined Phosphodiesterase PDE3/PDE4 Inhibition over PDE4 Inhibition Alone on Glucocorticoid- and Long-Acting 2-Adrenoceptor Agonist-Induced Gene Expression in Human Airway Epithelial Cells. Mol Pharmacol. 2014;

63. Pace E, Ferraro M, Uasuf CG, Giarratano A, Grutta S La, Liotta G, et al. Cilomilast counteracts the effects of cigarette smoke in airway epithelial cells. Cell Immunol. 2011;

64. Mackenzie SJ, Houslay MD. Action of rolipram on specific PDE4 cAMP phosphodiesterase isoforms and on the phosphorylation of cAMP-response-elementbinding protein (CREB) and p38 mitogen-activated protein (MAP) kinase in U937 monocytic cells. Biochem J. 2015;

65. Card GL, England BP, Suzuki Y, Fong D, Powell B, Lee B, et al. Structural basis for the activity of drugs that inhibit phosphodiesterases. Structure. 2004;

66. Giembycz MA. Cilomilast: a second generation phosphodiesterase 4 inhibitor for asthma and chronic obstructive pulmonary disease. Expert Opin Investig Drugs. 2001;

67. Lagente V, Martin-Chouly C, Boichot E, Martins MA, Silva PMR. Selective PDE4 inhibitors as potent anti-inflammatory drugs for the treatment of airway diseases. In: Memorias do Institute Oswaldo Cruz. 2005.

